# Diverse methanotrophs from alpha and gammaproteobacteria dwell in a stone quarry: a study from Western India wetland

**DOI:** 10.1101/2023.01.11.523701

**Authors:** Jyoti A. Mohite, Shubha S. Manvi, Shrinidhi Deshpande, Sanjana Patange, Rahul A. Bahulikar, Kajal Pardhi, Monali C. Rahalkar

**Author notes:** Accession numbers: *pmoA*: OQ354217, OQ354219, OQ354221, 16S rRNA gene: OQ373008, OQ373314, OQ373316, OQ373317.

## Abstract

Freshwater wetlands are a rich habitat for aerobic or micro-aerophilic methanotrophs. In the present study, we sampled a wetland ecosystem from Western India, a stone quarry with water situated amidst a hill. The wetland is surrounded by the macrophyte *Typha* sp. (cat-tail). To study the methanotroph diversity in this ecosystem, we collected mud samples from the mini-lakes and rhizospheric soil of *Typha* by random sampling. A serial dilution enrichment in liquid followed by isolation on agarose plates in a methane-air atmosphere was used. The high abundance of methanotrophs was indicated by the fact that growth was obtained till dilutions: 10^−9^ or 10^−10^. In *Typha* root-associated soil, Type I methanotrophs: *Methylomonas koyamae* and uncultured *Methylococcus* were isolated from the lower dilutions: 10^−3^-10^−6^. Type II methanotrophs *Methylocystis sp*. and *Methylosinus* spp were isolated from the higher dilutions. The *Methylococcus* strain isolated was about 97.4% (16S rRNA gene similarity) related to *Methylococcus capsulatus* and *Methylococcus geothermalis* and was predicted to represent a putative novel species. Three more methanotrophs, non-axenic, related to *Methylomagnum-Methylocaldum spp, Methylocucumis oryzae*, and *Methylocystis* spp, were cultured. Their affiliation was predicted based on the *pmoA* gene sequence. The mud samples showed relatively less diversity and mainly yielded two *Methylomonas* strains related to *M. koyamae*.

---

While various natural and anthropogenic activities emit methane, wetlands account for massive emitters and absorbers of methane gas [1]. Wetlands occupy 3.8% of the Earth’s land surface, amounting to about 20–40% of global CH_4_ emissions. Wetlands have anoxic zones where methane is produced. Methanotrophs utilize over 18 to 90% of the methane in wetlands [1], thus simultaneously establishing an equilibrium between methanogenesis and methanotrophy. [1]. Methanotrophs are aerobic to micro-aerophilic bacteria that use methane as the sole carbon source for growth and energy [2]. Aerobic methanotrophs associated with the roots of emergent aquatic plants are the primary colonizers of aquatic environments by making carbon available via methane consumption. Methane consumption by methane-oxidizing bacteria decreases the amount of methane that could be emitted by vegetation [3]. Methane-oxidizing bacteria in the oxic layer or the soil’s oxic-anoxic interphase carry out methane oxidation. Arctic, Antarctic, temperate wetlands, and lakes have been studied in detail [4-7], but tropical wetlands have been less studied for their methanotroph ecology.

Stone quarries are unique ecosystems, and the lakes created naturally in such quarries become a unique habitat for methanotrophs. In the present study, we sampled a wetland ecosystem associated with a stone quarry situated amidst hills in Pune city (known as ARAI hills), one of the only natural green habitats in the city. Due to the proximity of the institute ARAI (Automotive research association of India), the hill is popularly known as ARAI hill and is renowned as a potential niche for diverse flora and fauna [8]. The water-filled quarry contains 5-6 mini-lakes (Figure 1) and is rich in vegetation with aquatic weeds (*Typha* spp) and small water plants. One such aquatic weed is *Typha* spp, dominant in the west to the southwestern region of India. Most such investigations have been done on a few plants like rice and *Sphagnum* moss; however, various studies on the interaction between methanotrophs and plants will deepen our understanding of the carbon cycle and help regulate it [9]. Two earlier reports indicated that *Typha* species inhabit methanotrophs in their roots and rhizospheres [1,10]. Almost no data is available on the methanotrophs dwelling in Indian freshwater wetlands apart from our study where a novel genus and species from Type Ib methanotrophs was cultured from a brackish water wetland near a beach, *Methylolobus aquaticus* [11] and *Ca*. ‘Methylobacter oryzae’ from a rice field near Allepey wetland in Kerala [12]. The present study is one of the first attempts to document the methanotroph diversity in another freshwater ecosystem using a cultivation method proven to culture a large diverse population [13].

**Figure no. 1:**
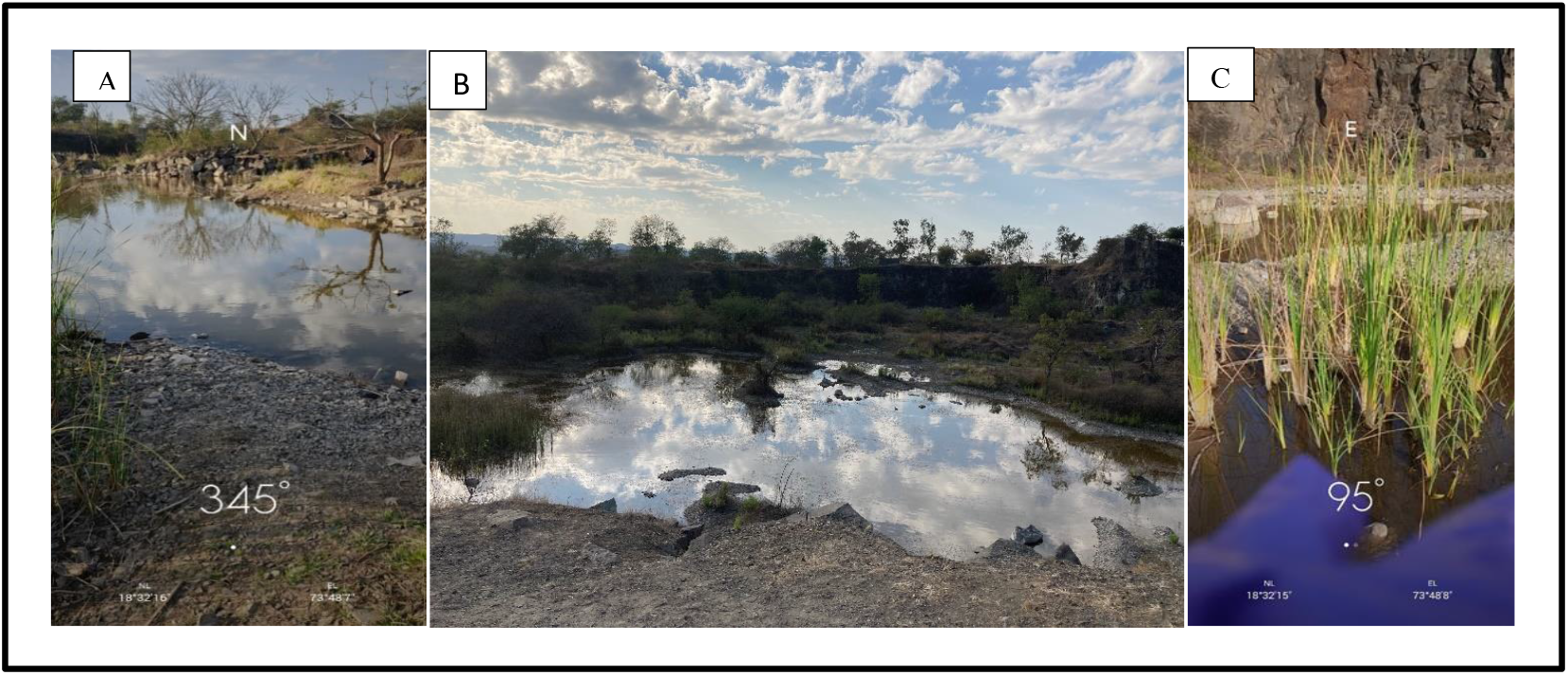
Site of sample collection: 18°32’ N, 73° 48’ ARAI quarry, Pune a, b and c. *Typha* plants

Mud samples and soil adhered to *Typha* root samples were collected from the wetland patch in Pune, India 18°32’ N, 73° 48’E (Figure 1) on 30 January 2022 (one root soil sample) and 16 February 2022 (three rhizosphere samples and three mud samples), Figure 1. All the samples were collected using gloves in sterile plastic vials or sterile plastic bags and proceeded for enrichment and oxidation of methane. Roots of *Typha* with the attached soil were collected by uprooting the plant. The roots were briefly washed in distilled water, and only the adhered soil with cut roots was used as a sample. Mud samples were collected from three different mini-lakes present at this site using sampling vials (50 ml capacity) from the sides of the lakes with about 10-15 cm water layer on the top.

Initially, we saw growth of methanotrophs in a serial dilution series set up with a single root sample of a *Typha* collected on 30 January 2022, and growth was seen up to 10-6 dilutions within two weeks of incubation. Thus, this site was found to be promising for the enrichment and isolation of methanotrophs. Therefore, the second sampling was conducted on 16 February 2022 to study the overall methanotroph diversity in this area. Serial dilutions were set up using modified Nitrate Mineral Salts referred to as dilute NMS medium was prepared [14] from 10^−1^ to 10^−10^ by adding 1g of the sample to a 9 ml sterile (dilute NMS) medium in 35 ml serum bottles [14]. A headspace volume of 20% was removed with a sterile syringe and filled with 20% methane. All the further procedures were done as described in our method article [13], except we directly proceeded for isolation after the first serial dilution enrichment step, as relatively high number of methanotroph like cells were seen. In all the samples collected from the quarry, growth was obtained till 10^−9^ or 10^−10^ dilutions in all the samples (Figure 2), indicating the high abundance of methanotrophs (∼10^−9^-10^−10^ cells/ g fresh weight of methanotrophs). The presence of abundant methanotrophs in our study site indicates that an active methane cycle is present here. Microscopically, methanotrophs were either in the form of large cells, oval or slightly elongated with dark interiors were seen in higher dilutions (indicative of Type I methanotrophs) or small coccoid cells (indicative of Type II methanotrophs) were observed in higher enrichment dilutions of 10^−9^ and 10^−10^ (Figure 2). Five pure cultures of methanotrophs named as strains AQ1-AQ5 were obtained in all from various dilutions (Figure 2, Table 1). DNA extracted from these pure cultures was subjected to 16S rRNA gene and *pmoA* gene sequencing. Three more cultures were obtained, showing a minor amount of small-sized bacteria which could be heterotrophs associated with the methanotrophs. A single morphotype with large cells of > 5µm length and oval shape indicative of Type I methanotrophs was seen in these cultures, indicative of *Methylomagnum* and *Methylocucumis*. We characterized the methanotroph present in these cultures (culture AQ6 and AQ7) using DNA extraction followed by *pmoA* PCR and sequencing and confirmed the predicted results obtained by morphological examination (Table 2). Both *Methylomagnum* [15]and *Methylocucumis* [16, 17] are known to be methanotrophs with large cells (∼5µm) with dark inclusions, and hence characteristic in morphology. The third culture (AQ8) showed small coccoid cells was indicative of Type II methanotrophs, identified as *Methylocystis*.

**Table no. 1:** Summary of type I and type II methanotrophs obtained in this study.

| Sr No. | Name of the methanotroph at genus level & NCBI accession number | Sample name & positive dilutions<br>Sampling date: 30.01.2022 (1)<br>Sampling date: 16.02.2022 (2) | Universal 16s rRNA gene sequence<br>BLAST hit name, percent similarity and NCBI accession number | *pmoA* gene sequence BLAST hit name, percent similarity and NCBI accession number |
| --- | --- | --- | --- | --- |
| 1. | *Methylosinus* sp. AQ1<br>16s (OQ373008) | ARAI root (1) sample dilution 10<sup>-6</sup><br>(2) | *Methylosinus sporium* 98.99%<br>(NR026512.1) | *Methylosinus trichosporium* 94.55%<br>(CP023737.1) |
| 2. | *Methylomonas* sp. AQ2<br>16s (OQ373314) | ARAI root (1) sample dilution 10<sup>-3</sup><br>(2) | *Methylomonas koyamae* 99.27%<br>(AP019777.1) | *Methylomonas koyamae* 96.06%<br>(AP019777.1) |
| 3. | *Methylomonas* sp. AQ3<br>16s (OQ373316) | ARAI mud (7) sample dilution 10<sup>-5</sup><br>(2) | *Methylomonas koyamae* 99.27%<br>(AP019777.1) | *Methylomonas koyamae* 96.29%<br>(AP019777.1) |
| 4. | *Methylococcus* sp. AQ4<br>16s (OQ373317) | ARAI root (1) sample dilution 10<sup>-6</sup><br>(2) | *Methylococcus capsulatus* 97.47%<br>(NR042183.1) | *Methylococcus geothermalis* 94.58%<br>(CP046565.1) |
| 5. | *Methylomonas* sp. AQ5<br>16s (OQ373316) | ARAI mud (7) sample dilution 10<sup>-9</sup><br>(2) | *Methylomonas koyamae* 99.27%<br>(AP019777.1) | *Methylomonas koyamae* 96.29%<br>(AP019777.1) |
| 6. | *Methylocucumis* sp. AQ6<br>*pmoA* (OQ354217) | ARAI root sample dilution 10<sup>-4</sup><br>(1) | Non-axenic culture | *Methylocucumis oryzae* Sn10-6 97.93%<br>(KT180167.1) |
| 7. | *Methylomagnum* sp. AQ7<br>*pmoA* (OQ354221) | ARAI water sample dilution 10<sup>-3</sup><br>(1) | Non-axenic culture | *Methylomagnum ishizawai* 89.45%<br>(AP019783.1) |
| 8. | *Methylocystis* sp. AQ8<br>*pmoA* (OQ354219) | ARAI root sample dilution 10<sup>-9</sup><br>(1) | Non-axenic culture | *Methylocystis hirsuta* CSC1 97.07%<br>(DQ364434.1) |

**Figure no. 2:**
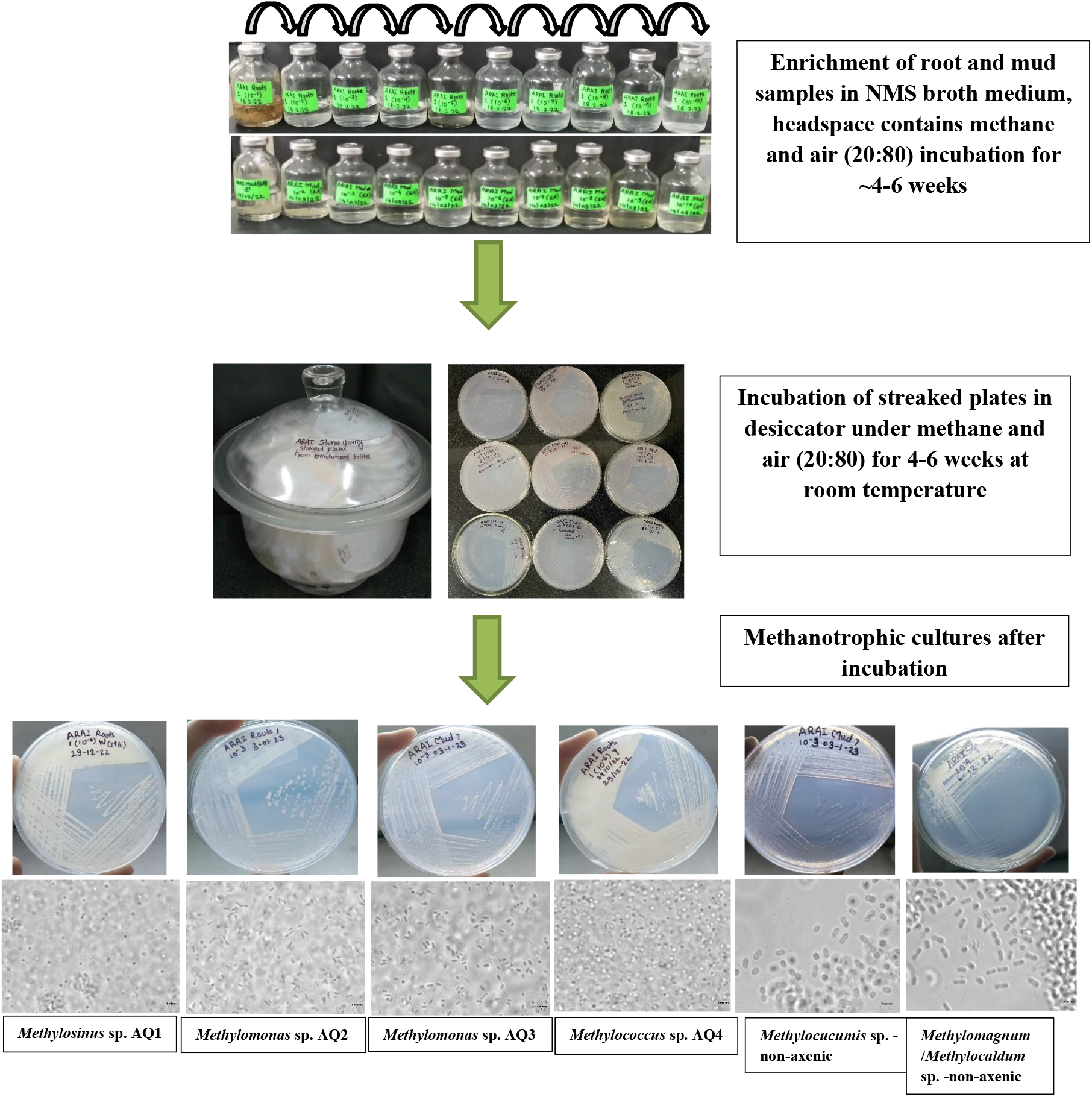
Enrichment, isolation, and identification of methanotrophs from ARAI stone quarry *Typha* plant

Thus, from *Typha* root-associated soil, Type I methanotrophs from the genera: *Methylomonas, Methylococcus, Methylomagnum* and *Methylocucumis* were isolated/ cultured from the lower dilutions: 10^−3^-10^−6^. Type II methanotrophs *Methylocystis, Methylosinus* spp were isolated/cultured from the higher dilutions (10^−9^), Table 1. The mud samples yielded two *Methylomonas* strains related to *M. koyamae*, Table 1. The *Methylococcus* strain isolated from 10^−6^ dilution was 97.4% related to *Methylococcus capsulatus and Methylococcus geothermalis*, on the basis of 16S rRNA gene and the partial *pmoA* gene was 95% related to that of *Methylococcus geothermalis* and the culture could represent a putative novel species. We found that small coccoid cells representing Type II methanotrophs of the Methylocystis-genus producing even milky, white turbidity were seen in all of the higher dilutions higher dilutions-10^−8^,10^−9^-10^−10^ in the root-associated methanotroph community. This was confirmed by further isolating *Methylocystis* from 10^−9^ dilution. On the contrary, Type I methanotrophs representing *Methylomonas, Methylocaldum-Methylomagnum and Methylocucumis* were present in lower dilutions, strengthening the observations made earlier in a Chinese wetland study [1]. In a Chinese wetland, it was observed that *Methylomonas* and *Methylocystis* dominated the root-associated community of *Typha* species, and the active community was composed of *Methylocystis* mainly [1]. *Typha* is a popular aquatic weed primarily occurs in tropical parts of the world. Strong associations of methanotrophs to the roots of emergent weeds have been published previously [10]. Further studies focusing on the methanotrophs present associated with *Typha* species in wetlands of various kinds would be more useful. The isolation of *Methylococcus* spp related to known species on a 97.4% similarity level indicates that this habitat may have yet un-cultured methanotrophs and could be a rich source for further isolations. Though *Methylococcus capsulatus* is the most studied methanotroph, *Methylococcus* genus has only two valid species, the classical methanotroph *Methylococcus capsulatus* [18] and the recently described *Methylococcus geothermalis* [19], and hence this niche could be an exciting habitat for the isolation of new species.

In Pune, temperatures can rise up to 42-43°C in summer, and the temperature of the water can increase in summer to a considerable extent. In summers, the wetland part also gets dried up, and hence we found that the roots were dominated by the presence of heat and desiccation-resistant methanotroph species: *Methylocystis, Methylosinus*, and *Methylococcus* [18, 19]. Annual monitoring of this habitat would give us a better picture of temporal diversity of methanotrophs.

The environmental relevance of methanotrophs as well as their potential applications in methane mitigation, are both equally important, and hence cultivation of methanotrophs becomes important to get deeper insight [20]. In the present study, we were successful in culturing important members of methanotrophs from a crucial ecological niche, a freshwater wetland, and methanotrophs associated with *Typha* spp. This study would lay a foundation for further studies focusing on methane mitigation from such sites.

## AUTHOR CONTRIBUTIONS

MR designed the experiments, collected the samples, guided all the experiments, and wrote the manuscript. RB, JM, SM, SD, and SP did the sampling and performed the enrichment and isolation of the methanotrophs. JM performed the phylogenetic analysis and prepared the trees. KP assisted in the isolation and maintenance of cultures. RB and MR found this unique sampling site, did the initial sampling, and co-designed the experiments. RB, JM, and SM assisted in writing the manuscript. All the authors reviewed the manuscript and approved the final version.

## ACKNOWLEDGMENTS

MCR acknowledges SERB for the POWER fellowship (SPF/2022/000045) provided to her. KP is thankful to UGC for JRF and JAM to SARTHI for providing the JRF. SSM acknowledges SERB project: CRG/2021/000941 for the JRF.

## Notes

### Competing Interest Statement

The authors have declared no competing interest.

### Summary of Updates

Figure 1, 2 and Table added. Isolation of methanotroph pure cultures added. Accession numbers added. Table and figures updated.

## References

1. Cui J, Zhao J, Wang Z, Cao W, Zhang S, Liu J, Bao Z (2020) Diversity of active root-associated methanotrophs of three emergent plants in a eutrophic wetland in northerm China AMB Express 10: 1–9.

2. Hanson RS, Hanson TE (1996) Methanotrophic bacteria. Microbiology and Molecular Biology Reviews 60: 439–471.

3. Laanbroek HJ (2010) Methane emission from natural wetlands: interplay between emergent macrophytes and soil microbial processes. A mini-review. Annals of Botany 105: 141–153.

4. Graef C, Hestnes AG, Svenning MM, Frenzel P (2011) The active methanotrophic community in a wetland from the High Arctic. Environmental microbiology reports 3: 466–472.

5. Michaud AB, Dore JE, Achberger AM, Christner BC, Mitchell AC, Skidmore ML, Vick-Majors TJ, Priscu JC (2017) Microbial oxidation as a methane sink beneath the West Antarctic Ice Sheet. Nature Geoscience 10: 582–585.

6. Wartiainen I, Hestnes AG, McDonald IR, Svenning MM (2006) Methylobacter tundripaludum sp. nov., a methane oxidizing bacterium from Arctic wetland soil on the Svalbard islands, Norway (786 N). International Journal of Systematic and Evolutionary Microbiology 56: 109–113.

7. Rahalkar M (2007) Aerobic methanotrophic bacterial communities from sediments of Lake Constance. University of Konstanz

8. Patil R, Mahajan M (2018) Herbaceous flora of weeds growing at ARAI hills. International journal of researches in biosciences, agriculture and technology: 16–19.

9. Yoshida N, Iguchi H, Yurimoto H, Murakami A, Sakai Y (2014) Aquatic plant surface as a niche for methanotrophs. Frontiers in Microbiology 5: 30. doi: 10.3389/fmicb.2014.00030

10. Fausser AC, Hoppert M, Walther P, Kazda M (2012) Roots of the wetland plants Typha latifolia and Phragmitis australis are inhabited by methanotrophic bacteria in biofilms. Flora 207: 775–782.

11. Rahalkar MC, Khatri K, Mohite J, Pandit P, Bahulikar R (2020) A novel Type I methanotroph Methylolobus aquaticus gen. nov. sp. nov. isolated from a tropical wetland. Antonie van Leeuwenhoek 113: 959–971.

12. Khatri K, Mohite JA, Pandit PS, Bahulikar RA, Rahalkar MC (2019) Description of ‘Ca. Methylobacter oryzae’ KRF1, a novel species from the environmentally important Methylobacter clade 2. Antonie van Leeuwenhoek 113: 729–735.

13. Rahalkar M, Khatri K, Pandit P, Bahulikar R, Mohite J (2021) Cultivation of Important Methanotrophs From Indian Rice Fields. Frontiers in Microbiology 12: 1–15.

14. Pandit PS, Rahalkar MC, Dhakephalkar PK, Ranade DR, Pore S, Arora P, Kapse N (2016) Deciphering community structure of methanotrophs dwelling in rice rhizospheres of an Indian rice field using cultivation and cultivation-independent approaches. Microbial Ecology 71: 634–644.

15. Khalifa A, Lee CG, Ogiso T, Ueno C, Dianou D, Demachi T, Katayama A, Asakawa S (2015) Methylomagnum ishizawai gen. nov., sp. nov., a mesophilic type I methanotroph isolated from rice rhizosphere. International Journal of Systematic and Evolutionary Microbiology 65: 3527–3534.

16. Pandit P, Rahalkar MC (2018) Renaming of ‘Candidatus Methylocucumis oryzae’ as Methylocucumis oryzae gen. nov., sp. nov., a novel Type I methanotroph isolated from India. Antonie Van Leeuwenhoek 112: 955–959.

17. Pandit PS, Hoppert M, Rahalkar MC (2018) Description of ‘Candidatus Methylocucumis oryzae’, a novel Type I methanotroph with large cells and pale pink colour, isolated from an Indian rice field. Antonie Van Leeuwenhoek 111: 2473–2484.

18. Bowman J (2000) The methanotrophs - the families Methylococcaceae and Methylocystaceae. In: Dworkin, M, Falkow, S, Rosenberg, E, Schleifer, K-H, Stackebrandt, E (eds.) The Prokaryotes: A handbook on the biology of bacteria: ecophysiology, isolation, identification, applications. Springer Verlag, New York

19. Awala SI, Bellosillo LA, Gwak J-H, Nguyen N-L, Kim S-J, Lee B-H, Rhee S-K (2020) Methylococcus geothermalis sp. nov., a methanotroph isolated from a geothermal field in the Republic of Korea. International Journal of Systematic and Evolutionary Microbiology 70: 5520–5530.

20. Cervantes F, Garcia S, Peura S, Balagurusamy N (2021) Editorial: Methanotrophs: Diversity, environmental relevance and applications. Frontiers in Microbiology 12:796861: 1–3.

